# A Statistical Detector for Ribosomal Frameshifts and Dual Encodings based on Ribosome Profiling

**DOI:** 10.1101/2022.06.06.495024

**Authors:** Alisa Yurovsky, Justin Gardin, Bruce Futcher, Steven Skiena

## Abstract

During protein synthesis, the ribosome shifts along the messenger RNA (mRNA) by exactly three nucleotides for each amino acid added to the protein being translated. However, in special cases, the sequence of the mRNA somehow induces the ribosome to shift forward by either two or four nucleotides. This shifts the “reading frame” in which the mRNA is translated, and gives rise to an otherwise unexpected protein. Such “programmed frameshifts” are well-known in viruses, including coronavirus, and a few cases of programmed frameshifting are also known in cellular genes. However, there is no good way, either experimental or informatic, to identify novel cases of programmed frameshifting. Thus it is possible that substantial numbers of cellular proteins generated by programmed frameshifting in human and other organisms remain unknown. Here, we build on prior work observing that data from ribosome profiling can be analyzed for anomalies in mRNA reading frame periodicity to identify putative programmed frameshifts. We develop a statistical framework to identify all likely (even for very low frameshifting rates) frameshift positions in a genome. We also develop a frameshift simulator for ribosome profiling data to verify our algorithm. We show high sensitivity of prediction on the simulated data, retrieving **97.4%** of the simulated frameshifts. Furthermore, our method found all three of the known yeast genes with programmed frameshifts. We list several hundred yeast genes that may contain +1 or −1 frameshifts. Our results suggest there could be a large number of un-annotated alternative proteins in the yeast genome generated by programmed frameshifting. This motivates further study and parallel investigations in the human genome. Frameshift Detector algorithms and instructions can be accessed in Github: https://github.com/ayurovsky/Frame-Shift-Detector.

## 1 INTRODUCTION

A ribosomal frameshift is a translation phenomenon where the ribosome switches from one mRNA reading frame to another, to produce a novel protein. The switch may involve only some percentage of ribosomes, in which case two proteins are produced, one in the original reading frame, and one in the shifted frame [1]. A frameshift happens when the ribosome, as it is decoding the messenger RNA (mRNA) codon by codon slips and continues decoding in one of the two other alternative reading frames. In a *+1 frameshift* (Figure 1, Panel A), the ribosome skips one nucleotide and continues translation in the second frame, and in a **-1** *frameshift* (Figure 1, Panel B), the ribosome shifts forward the normal three nucleotides, then backtracks by one nucleotide and continues translation in the third frame.

**Figure 1.**
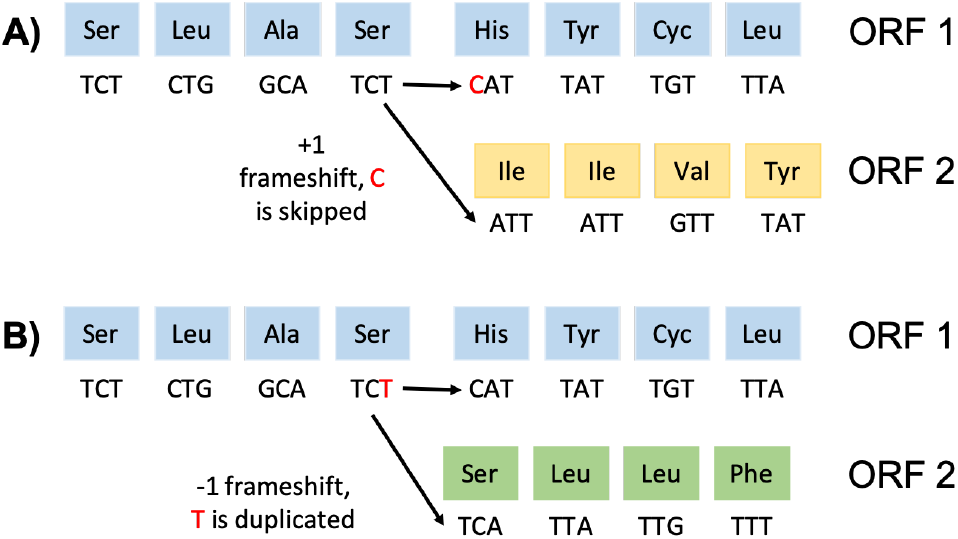
The most common types of frameshifts are the *+1 frameshift* (Panel A), where the ribosome skips one nucleotide and continues translation in the second frame, and the *-1 frameshift* (Panel B), where the ribosome backtracks by one nucleotide and continues translation in the third frame.

There are several proposed causes of ribosomal frameshifts including slippery sequences, RNA secondary structures, alternative splicing [2] and rare codons [3][4]. A slippery sequence is a small region on the mRNA that causes the ribosome to slip into the alternative coding frame. mRNA secondary structures such as stemloops and pseudoknots pause the ribosome, eventually making it slip into an alternative reading frame.

Known *-1 programmed frameshifts* typically require a slippery sequence, followed by a spacer region of 5-9 nucleotides, followed by a stem-loop RNA secondary structure [5][6]. The common slippery sequence for the programmed frameshift has a pattern of X XXY YYZ, where the Xs (usually) represent any three identical nucleotides, the Ys represent AAA or UUU, and Z stands for A, C, or U. While there exist examples of programmed *+1 frameshifts*[7], most *+1 frameshifts* do not require an RNA secondary structure, and their slippery sequences have a simpler format, comprising of codons that have rare transfer RNAs or out-of-frame stop codons [3][4][8][9][10].

There is increasing evidence that the rates of ribosomal frameshifting are controlled by trans-acting elements such as microRNAs, which can bind to RNA secondary structures and modulate their strength [11]. Other types of trans-acting molecules affecting frameshifting rates include amino-glycosides [12] and polyamines [13] and other proteins [5]. The frameshifting rates range from imperceptibly low rates to 100 percent in the case of yeast EST3 gene. In EST3, the ribosomes either fall off at the frameshifting site, resulting in a non-functional protein, or continue in the new frame, to produce a protein needed for telomerase function [9].

Ribosome frameshifting is ubiquitous in viruses, allowing them to encode more proteins in small genomes [14]. In contrast, *confirmed cases of frameshifting* are rare in bacterial and eukaryotic genomes. A few such cases have been very well studied, including the prfB gene of E. coli, encoding release factor 2; the EST3 and ABP140 genes of the yeast *S. cerevisiae,* encoding a component of telomerase, and a tRNA modification enzyme, respectively, and, most famously, the ornithine decarboxylase antizyme in many organisms [14]. The programmed frameshift to ornithine decarboxylase antizyme is highly conserved, probably because there is a regulatory mechanism by which polyamines (a product of ornithine decarboxylase) control the efficiency of frameshifting. That is, the programmed frameshift is part of the regulation of polyamine synthesis, and this is highly conserved [14].

While it is true that there are only a handful of known cases of programmed frameshifting in cellular organisms, it is also true that there are no systematic methods for identifying such cases. Thus it is possible that cellular organisms may contain significant numbers of proteins generated by programmed frameshifting of which we are still unaware. Indeed, Dinman and others have suggested that 1 to 10 percent of human and yeast genes could produce alternative protein products via frameshifting [14][15][16][17][18]. If true, this would substantially increase protein diversity in these organisms, and would likely involve new mechanisms of gene regulation. We thus have a complicated picture, where many eukaryotic genes could produce novel protein products at varying low rates due to endemic frameshifting, either accidental, or programmed.

Experimentally, information on the reading frame of ribosome translation can be gleaned from the method of Ribosome Profiling, which gives a high resolution snapshot of the locations of the actively translating ribosomes in a cell, by sequencing cDNA libraries generated from ribosome-protected fragments [19]. Because this method can give one nucleotide resolution, it can distinguish the reading frame in which any particular ribosome is probably translating. Usually, ribosomes are found translating in reading frame 1, which is the standard open reading frame. The general nature of our approach is that we analyze high-resolution ribosome profiling data sets; we assign probable reading frames to each ribosome-protected fragment; and we identify regions of mRNAs where “many” (in a statistical sense) ribosomes are in an alternative reading frame, either reading frame 2, or reading frame 3. In our informatic approach to frameshift detection, we focus on baker’s yeast — *Saccharomyces cerevisiae (S. cerevisiae),* a model organism that has been studied intensely by molecular genetics, including by ribosome profiling. This is in part due to the high quality of the existing ribosome profiling datasets for this organism.

More specifically, in this work, we develop a statistical framework to identify all likely frameshift positions in a genome, as well as a frameshift simulator for the ribosome profiling data to verify our algorithm.

Our main contributions are:

- **Statistical Framework** – We develop a framework to identify all likely (even for very low frameshifting rates) frameshift positions in a genome.
- **High Sensitivity** – We show high sensitivity of prediction on the simulated data, retrieving **97.4%** of frameshifts with low frameshifting rates.
- **Frameshift Detector Experimental Results** – We identify possible frameshift sites genome-wide for *S. cerevisiae*, with three known frameshifts being successfully assigned high scores by our algorithm.

The rest of the paper is organized as follows. In Section 2, we cover related work and the motivation for our study. In Section 3, we discuss the Bounded Probability Estimate – the central idea behind our statistical framework. In Section 4, we describe assignment of Ribosome Profiling reads to reading frames, the data. In Section 5 we detail our methodology for Frameshift Detector. In Section 6 we describe our frameshift simulator and the simulation results. In Section 7, we discuss the experimental findings. Section 8 concludes with lessons learned and future directions. Frameshift Detector algorithms and instructions can be accessed in Github: https://github.com/ayurovsky/Frame-Shift-Detector.

## 2 RELATED WORK AND PROBLEMS WITH STATE OF THE ART

Experimental discovery of new protein products due to frameshifting is a slow, expensive process that requires low-throughput, one- by one experimental methods. The three well known protein-coding genes in *S. cerevisiae* that undertake efficient +1 frameshifting are ABP140 (YOR239W), EST3 (YIL009C-A), OAZ1 (YPL052W). In these cases, the frameshifting was discovered by investigators who were, for various reasons, studying these particular genes [8][9][10]. The methods used would not have been appropriate for global, genomewide studies.

There also exist databases which compile experimentally validated and computationally predicted frameshift events from literature. In these cases, computational prediction involves various kinds of DNA sequence analysis, and so is limited to predicting events of kinds that are already understood. The Recode database summarizes up-to-date information from literature on verified frameshifts in a variety of organisms [20]. For *S. cerevisiae* there is a total of three frameshifting events out of about 6,000 protein coding genes, which underscores the difficulty in obtaining a validation set. The computational methods used by Recode to make novel frameshift predictions employ several narrow-domain Hidden Markov Models (HMMs) based on DNA sequence. ARFA [21] has trained an HMM to recognize and annotate *+1 frameshift* events in class-I bacterial release factors, RF1, RF2 and RFH, the bacterial genes responsible for proteins that terminate translation. OAF [7] has trained HMMs to recognize the *+1 frameshift* in antizymes – genes that regulate cellular polyamine levels and that are present in virtually all eukaryotes. Specific combinations of HMMs were trained for twelve groups that span major phyla.

Similarly to Recode, the FSDB database presents only the three experimentally validated frameshifting events in *S. cerevisiae* [22], as well as several individually-contributed un-validated predictions. FSDB also exposes the interface to its own computational frameshift search, FSFinder, which is a simple prediction of *-1 frameshifts* based on searching for slippery sequences (which includes many alternatives to the canonical sequences), followed by a spacer region, followed by a stem-loop RNA secondary structure inside of open reading frames. FSFinder predicts *+1* frameshifts for only two genes: the protein chain release factor and the antizyme (same as OAF), using slippery sequences that contain stop codons.

The PRFdb database contains only *-1 frameshift* predictions in eukaryotic genomes based on search for slippery sequences, followed by a spacer region, followed by RNA secondary structure [18]. The RNA secondary structure is re-confirmed using a variety of folding packages, and whole gene randomization was used to assess the statistical significance of the findings. The analysis yielded thousands of *-1 frameshift* predictions for *S. cerevisiae,* nine of which were selected for in-vivo experiments, yielding various percentages of frameshifting with a dual-luciferase frameshift reporter assay. We compare these to our findings in Section 7.

Another computational method, KnotInFrame, is more careful in selecting true pseudoknot secondary structures [23]. Both PRFdb and KnotInFrame focus on fairly stringent criteria for motif construction for predicting programmed *-1 frameshifts.*

For *+1 frameshift* prediction, FSCAN employs E. coli codon usage for tRNA competition between the first and second coding frames [4]. FSCAN lends more support for the idea that out-of-frame stops and in-frame rarely-used codons cause *+1 frameshifting;* we use this idea to obtain supporting evidence for our findings in Section 7. Another similar work demonstrated that CUU-AGG-C and CUU-AGU-U patterns, which are known to cause *+1 frameshifting* in ABP140 and EST3 genes, are highly depleted and predictive of *+1 frameshifting* in *S. cerevisiae* [24].

Finally, using a computational approach similar to Recode, GeneTack trains HMMs to distinguish overlapping ORFs that occur in the sequence due to a presence of a frame-shifted gene, from true overlapping or adjacent genes [25]. This approach identifies both *+1* and *-1* frame transitions, which can be due to programmed frameshifting events or alternative splicing, and was tested on frameshifts randomly inserted into genomes.

Again, we note that these computational methods are based on analysis of DNA sequence, and not on any experimental analysis of gene expression.

### 2.1 Ribosome Profiling Methods

Ribosome Profiling [19] is a scalable, high-throughput genomewide method for characterizing translation. In ribosome profiling, mRNAs are extracted from cells with ribosomes frozen onto these RNAs at exactly (ideally) the same positions as at the moment before extraction. The RNA protruding from each ribosome is removed by digestion with RNase, while the fragment of RNA over which the ribosome sits is protected from digestion. This fragment is about 28 nucleotides long. Many millions of these protected mRNA “footprints” are obtained in each experiment, and they are sequenced en masse, giving a record of the location of each ribosome on each mRNA in the cell population. The exact terminal nucleotides of the protected fragment are characteristic of a particular reading frame, and protected fragments from a single gene vary from one another by a multiple of three nucleotides, corresponding to the three-nucleotide shift of the ribosome during translation. Thus, the protected footprints report on the reading frame(s) of the ribosomes that generated the footprints. As we show here (e.g., Figure 2), the shift of ribosomes to an alternative reading frame can be detected by the characteristic sequences of the footprints.

**Figure 2:**
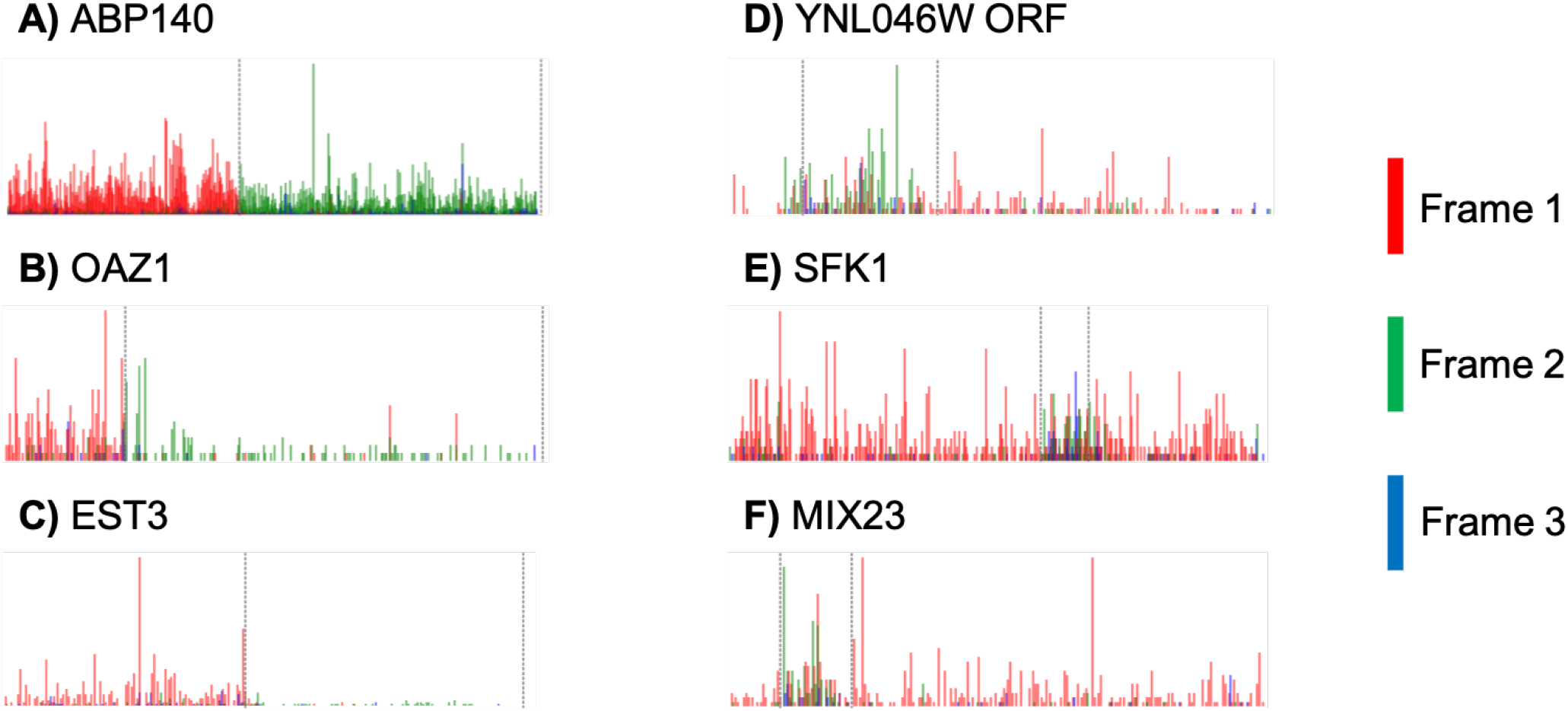
Ribosome profiling fragments mapped to six genes and assigned to reading frames by our method. The y-axis reflects the number of 5 ends mapping to each nucleotide position, and the color reflects the reading frame of those nucleotides. Panel A shows ABP140, a well known S. *cerevisiae* gene that undertakes efficient *+1 frameshifting.* The obvious red-to-green (frame 1 to frame 2) transition in the reading frame of the ribosome profiling fragments occurs exactly at the known position of the programmed frameshift. Panels B and C show OAZ1 and EST3, the other two genes known to undertake efficient *+1 frameshifting* in yeast. Panels D-F show three genes with potential partial frameshifts, evidenced by an increased number of fragments mapping to frames 2 or 3.

The challenges to use of ribosome profiling to determining reading frame are mainly that resolution and accuracy are not perfect; because of variation in the RNA digestion (and also in the conformation of the ribosome) there are a substantial number of protected RNA fragments that are slightly shorter or longer than the ideal. Furthermore, the vast majority of protected footprints are in reading frame 1, and, for most genes, the majority of footprints apparently in frames 2 or 3 are footprints actually from frame 1, where digestion has been slightly abnormal. Thus there is an issue of extracting a possibly small signal from noisy data, and a statistical approach is required.

Importantly, analysis of ribosome profiling datasets is an infor-matic method that is completely orthogonal to DNA-sequence based methods for detecting frameshifts (such as KnotInFrame, FSCAN, etc., see above). As such, it should be a powerful complement to such sequence-based approaches.

Genome-wide Ribosome Profiling [19] studies have become popular in the past decade, and a few have been used to verify or find frame shifts in prokaryotic and eukaryotic genomes. In recent works, ribosome profiling has been used in viruses and bacteria to verify or better characterize individual frameshifts. Ribosome profiling was used in encephalomyocarditis RNA virus to show that protein-directed frameshifting at a known site changes between 0 to 70 percent during the course of infection [5]. A specific cleaving technique has allowed better frame assignment in bacterial ribosome profiling and has verified a well-characterized *+1 frameshift* in prfB gene [26].

A Ribosome Profiling study on *S. cerevisiae* to quantify its behavior under hydrogen peroxide treatment used a simplistic heuristic for detecting frameshifts [27]. A small region after the stop codon of each gene was examined for footprint coverage similar to the gene; if footprints were found, a frameshift was called; several candidates were manually examined. Aside from OAZ1 and ABP140, this approach found four new *+1 frameshifts,* but only under the oxidative stress condition.

Ribosome Profiling has been used by Baranov and colleagues to detect frame transitions in the human genome, taking advantage of the triplet periodicity in the mapped ribosome protected fragments [17]. This approach assigns the fragments to frames disregarding the distributions of different fragment class lengths, as well omitting genes with relatively low expression (see Section 3). In this work, the cumulative subcodon proportion difference (CSCPD) is calculated for each frame for each codon, which is the difference between the proportion of fragments in the frame in a window before this position and after this position. Periodicity Transition Score (PTS) is calculated as the area of CSCPD excess over the 95th percentiles (using highly expressed genes) for each subcodon position. The PTS thresholds were then tuned to determine what should constitute a transition between frames.

### 2.2 Proteomic Methods

It is possible to verify putative protein products caused by frameshifts with proteomics. Using Mass Spectrometry (MS), peptides (cut in specific places from proteins) are detected by their specific masses. Then search algorithms, such as MASCOT, try to match these masses in databases for a given organism. There are several problems that must be overcome with adopting such an approach on a genome-wide level. First, because of how the peptides are generated by cutting, a careful identification of peptides that are only due to the frameshift must be made. Secondly, many frameshifts occur with low probabilities, and thus their potential protein products may not be picked up by MS. Thirdly, peptides not matched to known proteins are often discarded by studies, and appropriate datasets must be obtained or created that would have all of the recorded peptides [28].

Despite these problems, proteomics can provide evidence of novel proteins due to frameshifts. Several recent onco-proteogenomic studies used proteomic workflows to detect peptide variants within MS datasets to examine frameshift mutations in multiple human cancer cell lines [29][30]. A large-scale MS study of a model protozoan, Euplotes octocarinatus, demonstrated that 11.4% of genes require *+1 frameshifts* to produce complete gene products [31].

## 3 IDEA OF THE METHOD AND BOUNDED PROBABILITY ESTIMATE

We wanted to obtain a robust *p*-value for the presence of a frameshift (or dual-encoding for some other reason, see below). When a frameshift occurs, the number of fragments mapping to an alternative frame (2 or 3) increases, while the number of fragments mapping to the original frame (1) decreases. The percent change is reflective of the frameshifting efficiency – what proportion of the ribosomes shift (see Figure 2). For example (with a very high-quality dataset), before a frameshift, 95% of the fragments typically map to reading frame 1, with about 2.5% mapping to each of the other two frames. Assuming that this breakdown is about the same for every gene, imagine a frameshift in the middle of gene G. After the frameshift in G the local distribution of mapped fragments changes for the rest of this gene, where a much larger percentage (e.g., 50%) maps to (e.g.) frame 2, while 47.5% map to frame 1, and 2.5% to frame 3. The statistical significance of this shift in proportion will depend upon the efficiency of the shift, and the total number of protected fragments, which in turn depends on the level of gene expression, and the length of the shifted reading frame. Thus high statistical significance can be achieved by some combination of a highly efficient shift to the new reading frame, high gene expression (leading to a large number of fragments), or a long shifted open reading frame (again leading to a large number of fragments). (These are the same general considerations as in the work of Baranov and colleagues [17], but our specific mathematical approach is different.)

Having experimented with a number of statistical frameworks, we settled on using a binomial distribution to model the fragments mapping to frame 1 vs. frames 2 and 3. We obtain *q*, the probability of a fragment mapping to frame 1 by calculating the mean f1 proportion over all in frame nucleotide triplets in protein coding genes in a sample. Respectively, *p*, the probability of a fragment mapping to frame 2 or 3 is *p* = (1 – *q*). Then the probability of observing an event E, exactly k out of N fragments mapped to frames 2 and 3, in a given interval is:

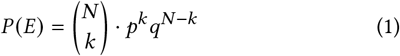

Probability of E is shown in Figure 3, Panel A. The ***p*-value** of a putative frameshift can then be expressed as the probability of observing k or more fragments mapped to frames 2 and 3, which is the sum over the tail of the binomial distribution:

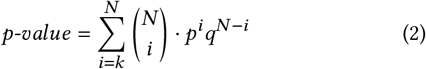

**Figure 3:**
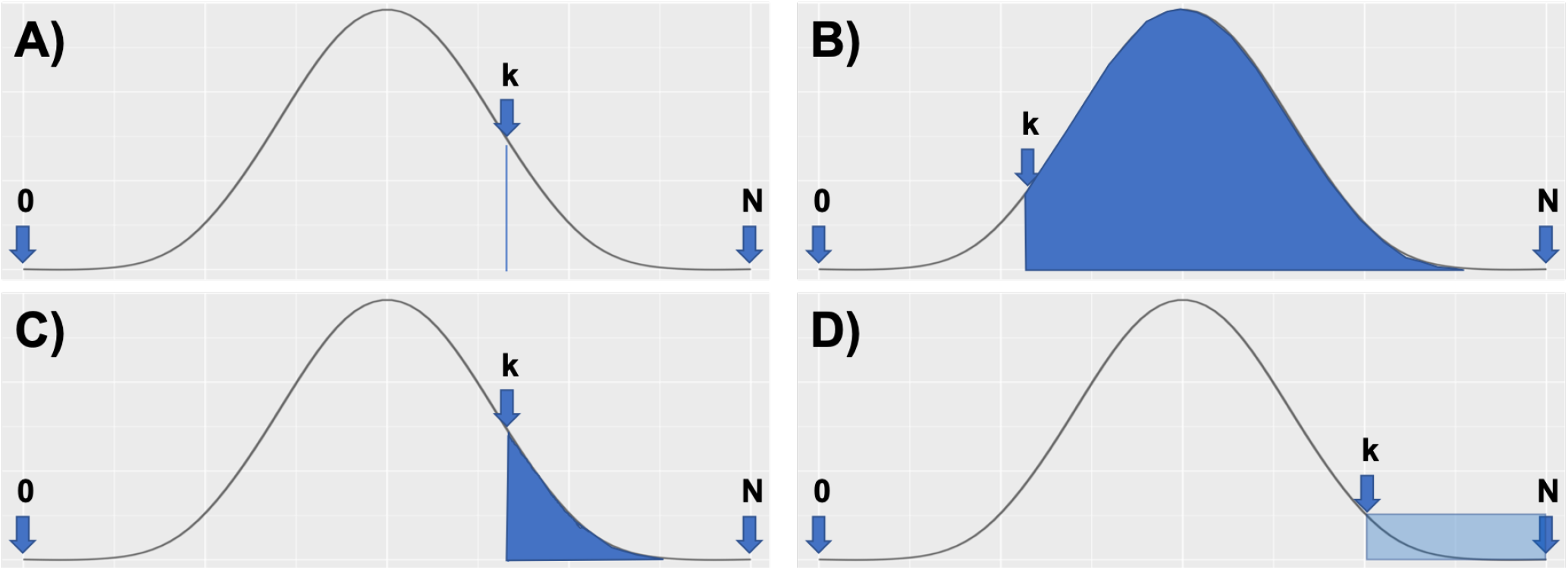
Visual representation of obtaining the p-value for a frameshift in a genomic interval with a total of N fragments, as described in Section 3. We use a binomial distribution to model the fragments mapping to frame 1 vs. frames 2 and 3. The blue line on Panel A shows the probability of observing an event where exactly k out of N fragments in a given interval are mapped to frames 2 and 3. The p-value of a putative frameshift can then be expressed as the probability of observing k or more fragments mapped to frames 2 and 3, which is the sum over the tail of the binomial distribution, shown as a blue shaded region in Panels B and C. The smaller proportion of f2 and f3 reads (a small value of k) will span the greater proportion of the total distribution, yielding high p-values (see Panel B), while observing a very high proportion of f2 and f3 reads (a large value of k) will yield highly significant p-values (see Panel C). Panel D shows that the upper bound on the probability estimate (the area of the colored rectangle) will yield significant p-values if the proportion is at the high tail of the distribution.

This makes intuitive sense, as the smaller proportion of f2 and f3 reads will span the greater proportion of the total distribution, yielding high *p*-values (see Figure 3, Panel B), while observing a very high proportion of f2 and f3 reads will yield highly significant *p*-values (see Figure 3, Panel C).

We have transformed the formula in several steps, due to the computational challenges presented by huge datasets. We took the logarithm of the binomial sum to prevent the overflow, to calculate the exponent of the *p*-value:

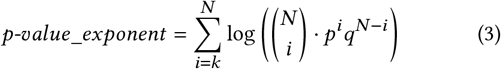

We then applied the Stirling Approximation to speed up the computation of the factorials in the sum:

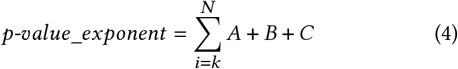

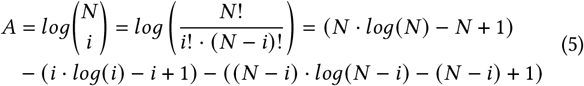

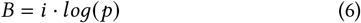

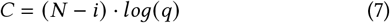

Finally, we observe that we can both obtain a more selective *p*-value and greatly speed up the computation by using an **upper bound** on the probability estimate (see Figure 3, Panel D). We replace the summation over the tail of the probability distribution in Equation 4 with multiplication by the length of the tail:

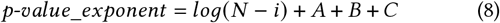

where *i* = *k*. The final run-time of genome wide frameshift detection algorithm using the bounded probability estimate on large(48 million uniquely mapped footprints) combined S. *cerevisiae* datasets is ten minutes on a single core.

## 4 FRAME ASSIGNMENT AND DATASET

### SELECTION

We use the standard processing of ribosome profiling data, obtaining the original SRA archives, clipping the adaptor sequences using the fastx toolkit, and aligning ribosome protected fragments to the *S. cerevisiae* genome using bowtie2. We only consider uniquely mapped reads.

Most good quality *S. cerevisiae* ribosome profiling datasets show a distribution centered on 28 nucleotide-long fragments, where the nucleotide triplet beginning with the 5’ nucleotide maps into frame 1. However, nuclease digestion is imperfect, and leads to a degree of noise. For instance, at the 5’ end, the nuclease might leave one extra nucleotide. In that case, one would have a 29 nucleotide fragment, with the 5’ nucleotide mapping to frame 3.

We have previously [32] dealt with noise and fragment length heterogeneity by grouping fragments into length classes (e.g., 27-mers, 28-mers, 29-mers, etc.), then, for each length class, determine how the majority of fragments in that class map to reading frame 1. For instance, for 29-mers, the triplet beginning at the second nucleotide typically maps to frame 3, so for all 29-mers, we assign a mapping based on the frame of the triplet beginning at the second nucleotide. This mapping can be experiment-specific. In some ribosome-profiling experiments, for some length classes, there might be significant heterogeneity in how fragments map to reading frames (e.g., a 70:30 split), and in such cases we would not use any fragments from this length class. Other experimental factors could also contribute to noise, and so could a certain low level of natural errors in maintaining the correct reading frame.

We have examined ten large recent ribosome profiling studies in *S. cerevisiae*. We omitted experiments designed to alter the normal behavior of *S. cerevisiae.* Some studies did not display a tight distribution centered on 28 nucleotide-long fragments, and/or did not have the vast majority of reads mapping to one frame per fragment length class, due to unknown experimental factors.

Ultimately, we selected the cleanest experiments from these four publications – Jan et. al. 2014 [33], Wu et. al. 2019 [34], Guydosh et. al. 2014 [35], and Young et. al. 2015 [36]. For each experiment, we only included the fragment length classes that were close to the peak of the length distribution and where at least 90% of the fragments mapped to a single frame.

The reading frame mapping of each length class was determined (see above), and the 5’ ends of individual fragments (adjusted +1 or −1 if necessary) assigned to genome positions. Thus, for each fragment we increment the counter for the genomic position of its 5’ end by one. These high quality assignments from the four experiments were then pooled into a single fragments file that we used for frameshift predictions. The fragments file contains the numbers of reads assigned to each nucleotide in each gene.

## 5 FRAMESHIFT DETECTOR

The inputs into the Frameshift detector are the fragments file, with the number of fragments mapped to each nucleotide in a gene, as described in Section 4, and *p*, the mean genome-wide probability of a fragment mapping to frame 2 or 3, as described in Section 3.

The core of the detector is a simple loop over all the genes, which finds the putative frameshift with the best *p*-value for each gene, as long as it is below the significance threshold. The significance threshold, for desired *α* = 0.05, was adjusted for multiple hypothesis testing with the Bonferroni Correction to be 10^-5^.

For each gene, we scan through each frame 2 and 3 position, and tabulate all fragments mapped to the interval from that position until the stop codon in the respective frame (2 or 3), if any, which we call the after_interval. We do the same for the same-length region upstream of the interval, called before_interval. Using the idea described in Section 3, we calculate the *p*-values for the putative frameshift in the before_interval and after_interval based on the proportion of reads mapping to frame 2 or 3.

To ensure a more stringent condition for the putative frameshifts, and to prioritize local change in translation over the change compared to the global average, we *subtract* the before_interval *p*-value from the after_interval *p*-value. This ensures that we only report putative frameshift regions that are not only highly significant, but are also highly significant relative to their local gene neighborhoods.

The intent of this method is to discover ribosomal frameshifts, but it will actually discover regions which are read by the ribosome in two (or three) reading frames for any reason. In addition to frameshifting, a gene might be read in another reading frame because there is an alternative Start codon either upstream or downstream of the annotated Start codon, and in a different reading frame. Simultaneous translation from two different Start codons in different reading frames can give overlapping regions with translation in two frames. Indeed, several of the genes found by Frameshift Detector appear to have dual encodings for this reason (see below).

## 6 SIMULATION RESULTS

Since the goal is to discover novel frameshifting sites, we needed an unbiased way of assessing the predictive power of the technique. We developed a simple simulation infrastructure which augments the ribosome fragments in frame 2 or 3 at random starting points in highly expressed genes, and applies the frameshift detector to the augmented dataset to test the recovery of inserted frameshifts with different percentage of augmentation and at different precision levels. The simulator selects highly expressed genes, with the average number of fragments mapped to a base of this gene being greater than 10. We randomly select the location for the start of the simulated shift (omitting the very beginning and end of the gene), and randomly select the direction of the shift – into frame 2 or 3; the shift information is saved. We then scan the genome sequence for the stop codon in the shifted frame, defining the region that will be augmented in the simulation. In the augmented region, we increase the number of reads in the shifted frame by the number of reads in frame 1 for the same codon, scaled by a range of multiplication factors. We create a separate augmented dataset for each multiplication factor.

We then process the augmented datasets with the Frameshift Detector and report number of recovered frameshifts at different levels of precision (see Figure 4). We can see that our method demonstrates reasonable sensitivity even at minuscule perturbations to the fragment allocation frequency among frames, calling 19.5% of the simulated frameshifts with the multiplication factor of 0.05 within 10 nucleotides from the actual simulated frameshift start, and 36.9% of those frameshifts when the precision criterion was relaxed to be within 30 nucleotides of the simulated start. We further note that at multiplication factor of just 0.3, the predictive ability reaches near maximum, recovering **86.2%** of the simulated frameshifts within 10 nucleotides from the actual simulated frameshift start, and **97.4%** of those frameshifts when the precision criterion was relaxed to be within 30 nucleotides of the simulated start.

**Figure 4:**
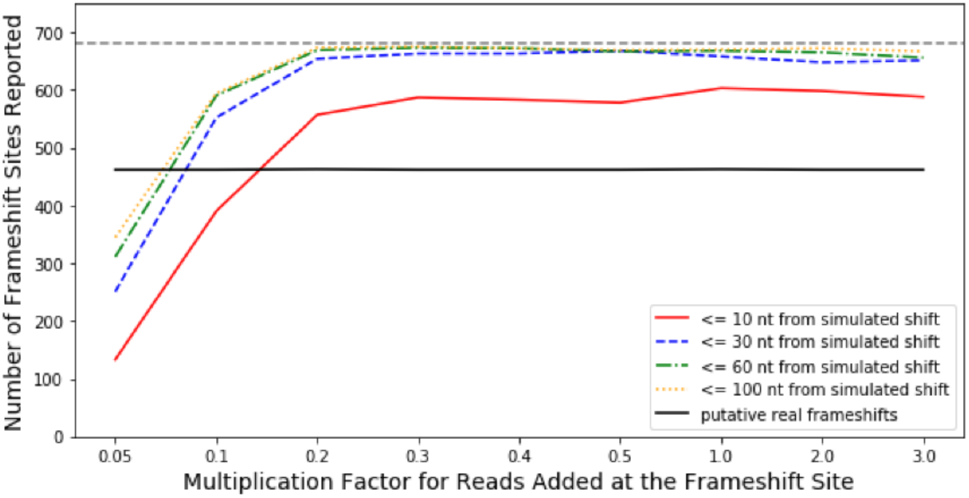
This figure demonstrates the ability of the frameshift detector to recover simulated frameshifts. The dashed line on the top shows the total number of simulated frameshifts. The colored lines show the number of recovered frameshifts at different precision thresholds for different read multiplication factors. The black line shows the (correctly constant) number of frameshifts found by the detector in genes not chosen for the simulation.

While our simulation results show high sensitivity, we are not able to assess the specificity of the Frame Shift detector, as there is a huge number of putative frame shift sites across the genome. Only three of these are known to be genuine frameshifts, and there is no clear way of validating or invalidating the novel sites. Close to five hundred frameshifts found by the detector in genes not chosen for the simulation (see Figure 4) are the frameshifts that our algorithm finds in poorly expressed genes. Full results are discussed in the following section.

## 7 FRAMESHIFT DISCOVERY RESULTS

Running Frameshift Detector on the combined *S. cerevisiae* Ribosome Profiling dataset (see Section 4) has yielded 831 candidate frameshifts which have passed the Bonferroni-corrected *p*-value threshold. Figure 5 shows the top 40 +1 frameshift results, while the top 572 are shown in Supplementary Table 1.

**Figure 5:**
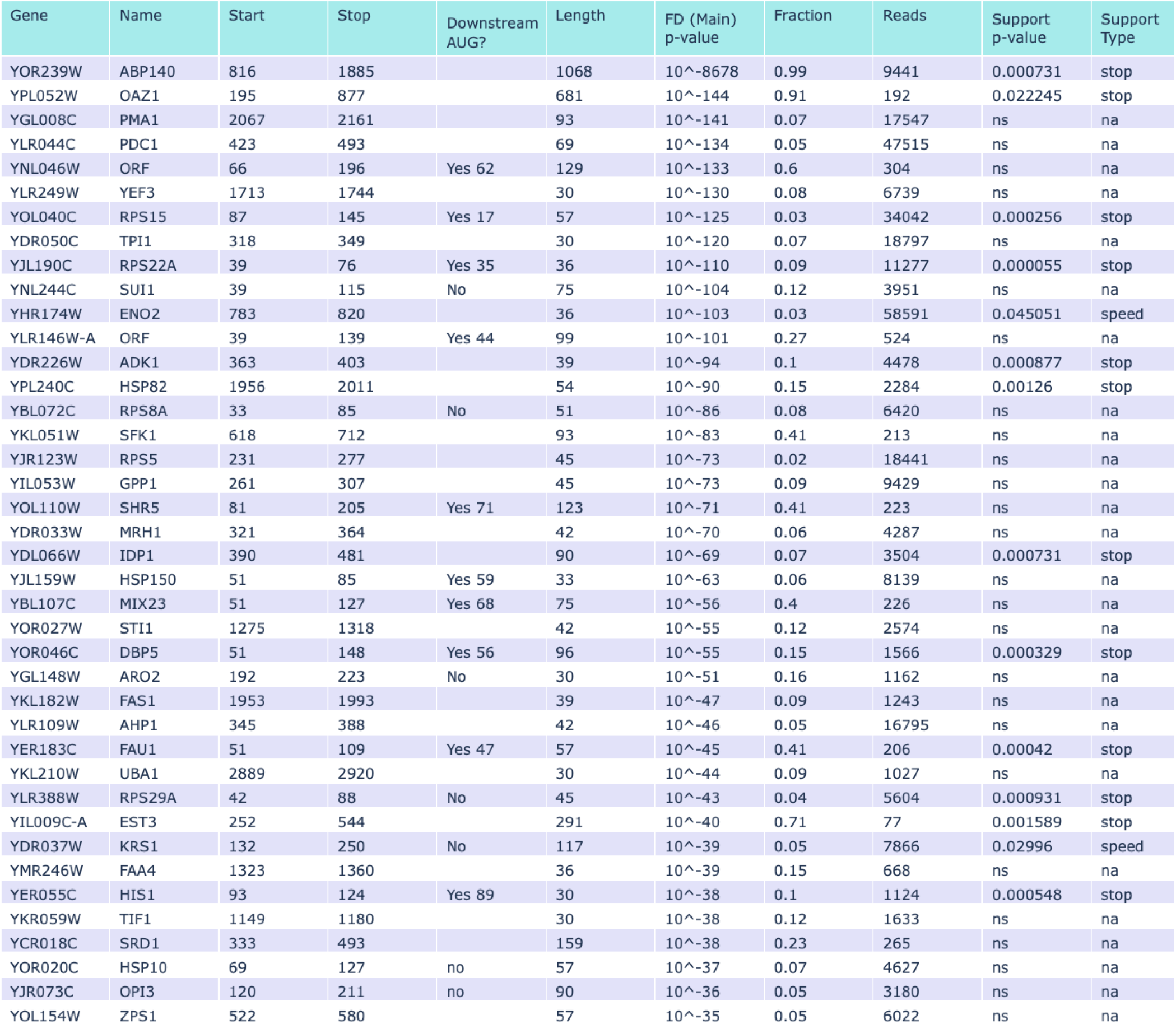
The top 40 results for the *+1 frameshifts,* sorted by the Frameshift Detector *p*-values (FD (Main) *p-value*). The Start and Stop positions give the location of the frameshifted region relative to the start of the gene. “Downstream AUG?” shows the nucleotide position (if present) of a downstream AUG suitable to explain the recoded region in selected genes. “Length” is the length (nucleotides) of the shifted region. Fraction is the proportion of reads in the shifted frame. The Reads column shows the number of ribosome fragments used in the calculation of the p-value. The final two columns show the p-value and type for the supporting signal (out-of-frame stop signal or the speed of codons before the frameshift, a proxy for rare transfer RNAs). Of the three well-established *+1 frameshifted* genes in *S. cerevisiae,* ABP140 (YOR239W) and OAZ1 (YPL052W) have the two best Frameshift Detector *p*-values, while EST3 (YIL009C-A) also appears in the top 40 results, with its less impressive *p*-value most likely due to lower coverage.

We first focus on the *+1 frameshifts.* In other systems, typically viruses, *+1 frameshifts* usually occur in the context of “slippery sequences”, which contain out-of-frame stop codons and/or stretches of codons with rare transfer RNAs. It was of interest to see whether the sites identified by Frameshift Detector were associated with these slippery sequences, as suggested by [24] and [4]. We analyze the sequences around the putative frameshift location as determined by Frameshift Detector for such slippery sequences, and compare the signal strength to a distribution from approximately 58 thousand random locations inside genes to obtain the statistical significance of the slippery sequence signal.

For stop codons, for each putative *+1 frameshift* location, we consider the distance from the closest out-of-frame stop codon, and calculate the *stop_p-value* as the rank in the distribution of such distances, calculated for randomly selected locations in protein coding genes. For the rare codon signal, we use the in-vivo codon translation speeds from Gardin et al. [32] as a proxy, as codon translation speeds show high correlation with rare transfer RNAs; slow speed of translation (while waiting for the transfer RNA) contributes to the ribosome slippage. We consider the total speed of the twenty codons around the putative frameshift, and calculate the *speed_p-value* as the rank in the distribution of such speeds, calculated for randomly selected locations in protein coding genes.

Figure 5 shows the top results for *+1 frameshifts,* sorted by the Frameshift Detector *p*-values and also showing *p*-values for the corresponding supporting signal. Most significantly, we note that the three well known protein-coding genes in *S. cerevisiae* that undertake efficient *+1 frameshifting,* ABP140 (YOR239W), OAZ1 (YPL052W) EST3 (YIL009C-A), are found in the top Frameshift Detector results.

Sixteen of the top 40 genes had an apparent frameshift within 150 nucleotides of the annotated Start codon. These seemed cases where dual-encoding might arise from usage of a second Start codon downstream of the annotated Start, rather than from a frameshift. We examined all 16 cases, and found that 10 of them (YNL046w, RPS15, RPS22a, YLR146W-A, SHR5, HSP150, MIX23, DBP5, FAU1 and HIS1 did indeed have an AUG triplet close to the observed beginning of the dual-encoded region, and in the correct frame for the stop codon ending the dual-encoded regions. Thus we believe these 10 dual-encoded regions arise from an alternative translational Start, and not from a frameshift.

The three longest shifted encodings in Figure 5 are for ABP140, OAZ1, and EST3, the three known cases of alternative encoding by frameshifting. All other shifted encodings are significantly shorter. We examined the five longest of these (KRS1, SRD1, SHR5, YLR146W-A, and YNL046W) using protein-protein Blast to see if the alternative (i.e., shifted) encodings matched any other protein in the database. None did.

Next, we look at the *-1 frameshifts*, where the signal is a slippery sequence followed by a spacer and then an RNA secondary structure. Figure 6 shows the top results for *-1 frameshifts*, sorted by the Frameshift Detector **p**-values. The top 161 results are shown in Supplementary Table 2. We attempted to obtain an orthogonal signal by comparing our results to the *-1 frameshifts* database PRFdb [18]. For each putative *-1 frameshift* location, we consider the distance from the closest PRFdb signal, and calculate the *struct_p-value* as the rank in the distribution of such distances, calculated for approximately 58 thousand randomly selected locations in protein coding genes. While some of *-1 frameshifts* have a statistically significant *struct_p-value,* none of the top results do.

**Figure 6:**
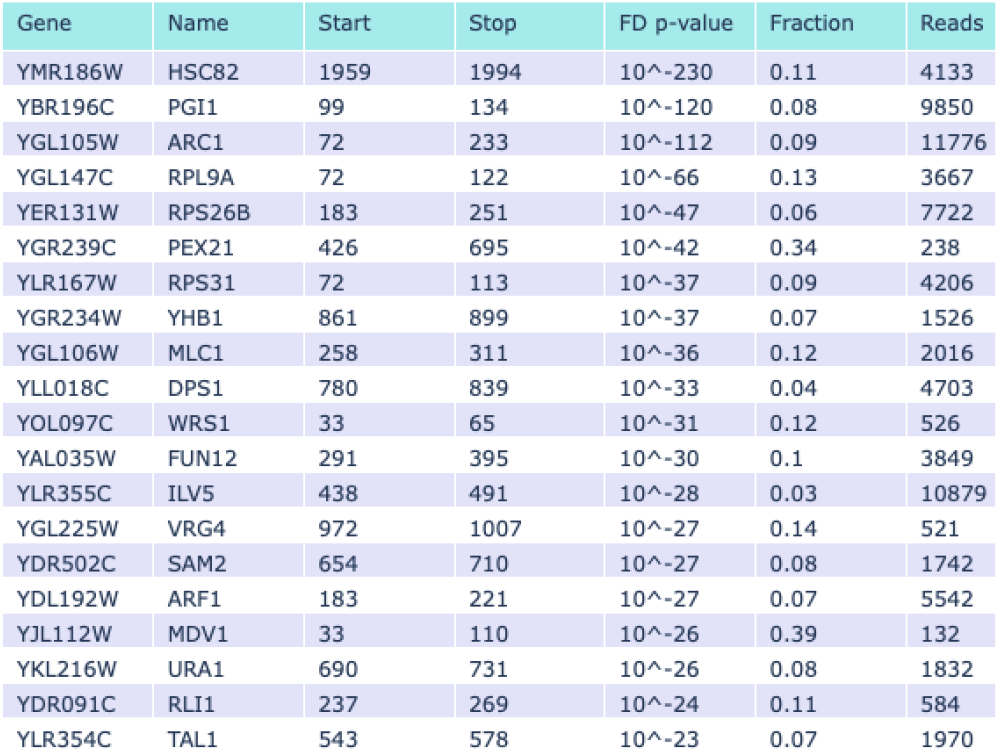
This figure shows the top 20 results for the *-1 frameshifts*, sorted by the Frameshift Detector p-values *p-value*). The Start and Stop positions give the location of the frameshifted region relative to the start of the gene. The Fraction is the proportion of reads in the shifted frame. The Reads column shows the number of ribosome fragments used in the calculation of the *p*-value.

We checked 6 genes PGI1, ARC1, RPL9a, RPS31, WRS1, and MDV1 for possible alternative AUG codons just inside the annotated open reading frame. Only ARC1 had an AUG suitable for explaining the observed region of dual-encoding. We also checked the longest regions of alternative encoding (from ARC1, PEX21, and FUN12) for similarity to other proteins in the database, but again found none.

We should note several things with regard to the found *-1 frame shifts*. First, we found few locations where our putative frameshifts are co-located with the PRFdb predictions, and the corresponding Frameshift Detector *p*-values for these locations are less significant than for our *+1 frameshift* findings. Secondly, when we further analyzed the genome around our most significant putative *-1 frameshift* sites, according to the Frameshift Detector *p*-values, we did not find any of the common *-1 frameshift* slippery sequences. This leads us to believe that either *S. cerevisiae* has novel sequence signals for *-1 frameshifting,* or there are few or no real *-1 frameshifts* in *S. cerevisiae*, or our methodology is imperfect for detecting *-1 frameshifts* in experimental data (despite showing high sensitivity in simulations).

Aside from *-1 frameshifts* and *+1 frameshifts* inside the body of the gene, we have identified around 100 genes where it appears that a certain fraction of ribosomes begin translation out of frame (relative to the gene) before the start codon. These are listed in supplementary Table 3. The translation initiation inside the 5’ UTR happens at out of frame ATG or alternative start codons, and has been found in both *+1* and *-1* frames.

Finally, for some genes such as YNL046W, SFK1, and MIX23 (Figure 2), the frameshifted region is a mixture of shifts to frame 2 and to frame 3. This could reflect some kind of confusion on the part of the ribosome, as opposed to a directed shift to one alternative frame.

## 8 CONCLUSION

We developed a statistical framework to identify all likely frameshift positions in a genome. We show high sensitivity of prediction on simulated data, retrieving 97.4% of frameshifts with low frameshifting rate. More importantly, Frameshift Detector is successful in finding known programmed frameshifts. In *S. cerevisiae*, there are three known cellular genes with a programmed frameshift: two of these, AMP140 and OAZ1, are the top two hits in Frameshift Detector, while the third, EST3, is the 32nd best hit. The 29 hits between OAZ1 and EST3 are good candidates for novel programmed frameshift genes. Beyond EST3, there are hundreds of candidate frameshifted genes with extremely small *p*-values. This motivates future studies to validate the candidates in Figure 5, and also to apply the same approach to other genomes, such as the human genome, to find still more potential programmed frameshifts and novel proteins.

AMP140, OAZ1, and EST3 stand out from other hits in Figure 5 in two ways. First, they have the highest fraction of ribosomes in the shifted frame (0.99, 0.91, 0.71, respectively), while the next highest shifted fractions, 0.60 (YNL046w), 0.41 (SFK1), 0.41 (SHR5) and 0.41 (Fau1) are noticeably smaller. Furthermore three of these “shifts” (YNL046w, SHR5, FAU1) are very close to the 5’ end of the gene, and easily explained by initiation at an alternative AUG codons in the shifted frame (see below). Second, they stand out because they have by far the largest length of shifted open coding region. We examined the novel proteins encoded by the eight next-longest shifted coding regions to see if they were similar to other known proteins, but none were. One possibility, therefore, is that *S. cerevisiae* actually has only three genes with adaptive (see below) programmed frameshifts (AMP140, OAZ1, and EST3), and Frameshift Detector has successfully found them all—that is, potentially, 100% sensitivity. In this case, one would want to apply Frameshift Detector to other genomes with suitable ribosome profiling data, in the hope that there too a large fraction of genuine, adaptive programmed frameshifts would be discovered.

In AMP140, OAZ1, and EST3, the programmed frameshift results in a significantly lengthened protein, where the additional protein sequence is important for function, and the incorporation of extra peptide sequence is the obvious function of the frameshifting. However, the opposite could also be adaptive—that is, in some circumstances, it might be adaptive for the cell to truncate a protein. A programmed frameshift might accomplish the goal of removing some C-terminal functional domain by frameshifting to a novel stop codon. One candidate for a frameshifted gene of this kind is SFK1. In this case, the frameshifted protein fairly quickly ends at a novel stop codon, and a part of the conserved Frag domain of Sfk1 is removed. Interestingly, Sfk1 has homologs in other yeasts where the region of Sfk1 before the frameshift is conserved, but the region after is not, suggesting that the pre-frameshift part of the protein has a function.

The intent of Frameshift Detector is to discover adaptive programmed frameshifts. However, none of the new regions discovered here show clear signs of being of this adaptive type. Operationally, Frameshift Detector finds regions where ribosomes are translating in unexpected reading frames. However, there are various kinds of events that can cause this beyond programmed frameshifting. First, there may be nucleotide sequences that are error-prone with regard to translation, and the ribosome may shift to a new frame (or frames) accidentally at a high rate. These frameshifts may be non-adaptive or mal-adaptive, but unavoidable. Identification of such sites may define what kinds of sequences are difficult for translation, and may inform as to error rates of translation. Second, ribosomes may be in unexpected reading frames because they have initiated from alternative, out-of-frame initiation codons. These could be either 5’ or 3’ of the annotated Start codon. Possible examples of this are YNL046w, SHR5, and FAU1. Third, defects in splicing (or alternative splicing) could lead to abnormal (or alternative) mRNAs with alternative Start codons, and these would appear to have out-of-frame translation. In our list, RPS29A and RPS8A both have 5’ UTR introns, and defective splicing coupled to an alternative Start codon could be seen by Frameshift Detector. Fourth, defective mRNAs made by insertion or deletion of a nucleotide at the transcriptional level would give out-of-frame translation. Finally, unknown and unidentified anti-sense mRNAs could be incorrectly seen as out-of-frame translation of the sense strand. All these events are of potential interest, and could be identified by Frameshift Detector.

## Supporting information

Supplemental Table 1 Plus One Shifts

Supplemental Table 2 Minus One Shifts

Supplemental Table 3 Dual Encodings from Upstream

## ACKNOWLEDGMENTS

This work was done with support from the NSF grant GRFP-1315232 and the NIH grant R01GM127542-04.

## Notes

### Competing Interest Statement

The authors have declared no competing interest.

